# Effects of MDMA on socioemotional feelings, authenticity, and autobiographical disclosure in healthy volunteers in a controlled setting

**DOI:** 10.1101/021329

**Authors:** Matthew J. Baggott, Jeremy R. Coyle, Jennifer D. Siegrist, Kathleen Garrison, Gantt Galloway, John E. Mendelson

**Keywords:** MDMA, ecstasy, entactogen, anxiety, authenticity, emotion, interpersonal, social

## Abstract

The drug 3,4-methylenedioxymethamphetamine (MDMA, “ecstasy”, “molly”) is a widely used illicit drug and experimental adjunct to psychotherapy. MDMA has unusual, poorly understood socioemotional effects, including feelings of interpersonal closeness and sociability. To better understand these effects, we conducted a within-subjects double-blind placebo controlled study of the effects of 1.5 mg/kg oral MDMA on social emotions and autobiographical disclosure in a controlled setting. MDMA displayed both sedative- and stimulant-like effects, including increased self-report anxiety. At the same time, MDMA positively altered evaluation of the self (i.e., increasing feelings of authenticity) while decreasing concerns about negative evaluation by others (i.e., decreasing social anxiety). Consistent with these feelings, MDMA increased how comfortable participants felt describing emotional memories. Overall, MDMA produced a prosocial syndrome that seemed to facilitate emotional disclosure and that appears consistent with the suggestion that it represents a novel pharmacological class.

## Introduction

The drug 3,4-methylenedioxymethamphetamine (MDMA, “ecstasy”, “molly”) is a widely used illicit drug and experimental adjunct to psychotherapy. MDMA is known among drug users for its socioemotional effects, such as feelings of empathy, interpersonal closeness, and sociability (Bravo, 2001; Kelly et al., 2006; Rodgers et al., 2006; Sumnall et al., 2006). Before it was classified as a controlled substance in the USA, MDMA was used as an adjunct to psychotherapy by therapists because it appeared to decrease defensiveness and enhance feelings of emotional closeness (Wolfson, 1986; Greer and Tolbert, 1986). More recently, clinical trials have tested MDMA as a therapeutic adjunct in patients with post-traumatic stress disorder (Mithoefer et al., 2013; Oehen et al., 2013). Thus, anecdotal and experimental data indicate that MDMA produces potentially therapeutic acute socioemotional effects.

There is not yet a mature scientific understanding of these acute socioemotional effects. One potential psychological mechanism of MDMA is that it may lessen sensitivity to threatening stimuli (Bedi et al., 2010; Bedi et al., 2009). In an early report, the clinicians Greer and Tolbert (1986) observed that MDMA lessened concern about threats, allowing events and ideas that were normally distressing to be addressed with reduced discomfort in psychotherapy. Consistent with this observation, some studies have reported MDMA may decrease ability to identify emotionally negative expressions, including fear (Hysek et al., 2012; Bedi et al., 2010), and may create a bias to identify expressions as emotionally positive (Hysek et al., 2012). These findings are consistent with decreased threat sensitivity, although they may partly result from a mood congruent bias in either response or perception.

The hypothesis that MDMA decreases threat sensitivity appears to be contradicted by findings that MDMA sometimes acutely increases rather than decreases anxiety. For example, Bedi and de Wit (2011) found that MDMA dose-dependently increased self-report VAS anxiety. This effect persisted when data from multiple studies in that and two other laboratories were pooled into a larger analysis (Kirkpatrick et al., 2014). MDMA did significantly decrease STAI-S anxiety scores at a late time point in one study (Liechti and Vollenweider, 2000) but no effect was seen in others from the same laboratory (Hasler et al., 2009; Liechti et al., 2000). In a pooled analysis of early studies from that group, MDMA increased apprehension-anxiety scores in females but not males (Liechti et al 2001). Overall, there is little evidence that MDMA has consistent clinically meaningful effects on anxiety.

We sought to clarify MDMA’s self-report effects on anxiety and affective processing. We hypothesized that MDMA might specifically decrease social anxiety. Social anxiety, or fear of negative evaluation, is considered a fundamental fear that is distinct from injury/illness sensitivity and anxiety sensitivity (the tendency to appraise anxiety-related cognitive changes and sensations as harmful) (Reiss, 1991; Taylor, 1993). Social anxiety seemed a plausible domain to measure since MDMA increases self-report sociability (suggesting decreased social anxiety) and this has been observed even when there were simultaneous increases in self-report anxiety (e.g., Bedi and de Wit, 2011). To measure general anxiety, we used a single VAS item because, as discussed above, longer validated scales have not shown consistent effects from MDMA.

Another potential psychological mechanism of MDMA is that it may increase sociability and alter appraisal of others. Although it may seem superficially contradictory to hypothesize increased sociability existing with increased anxiety, there is no actual contradiction. For example, concern about threats could trigger a protective sociality, as in the tend-and-befriend model of stress response (Taylor, 2006).

MDMA-induced self-report sociality is well demonstrated. Participants often report feeling increased closeness to others, kindness, or friendliness. There are also inconsistent reports of possible changes in evaluation of social stimuli (Wardle et al., 2014; Hysek et al., 2013). Wardle and de Wit (2014) found MDMA slightly but significantly increased ratings of perceived partner empathy, though it is unclear if the magnitude of the effect was clinically meaningful. In a therapeutic setting, social effects of MDMA may enhance the therapeutic alliance and, in couples therapy, may facilitate meaningful interactions.

Research on MDMA effects on sociability has focused on appraisal of others and little is known about how MDMA might alter self-appraisals. In addition to measuring concerns about negative appraisals from others (social anxiety), we therefore sought to measure changes in self-appraisal. We did this using the construct of authenticity, which can be thought of as knowing one’s thoughts and feelings and acting in accordance with them (Sheldon et al., 1997; Rogers, 1961; Goldman and Kernis, 2002). We selected this construct because several psychotherapists administering MDMA to patients had emphasized seemingly related effects. For example, Greer and Tolbert (1990) wrote that MDMA helped individuals to “experience their true nature,” while Adamson and Metzner (1988) hypothesized MDMA improved access to “one’s true self.” Self-report authenticity is associated with decreased defensiveness (Lakey et al., 2008) and increased well-being (Rogers, 1961; Maslow, 1968; Wood et al., 2008).

We collected self-report measures of anxiety, sociability, and authenticity in the context of an autobiographical speech task. This provided a consistent structured social experience that facilitated participant ratings of social functioning and it allowed us to examine whether MDMA altered remembering and describing of positive and negative psychological material. There have been several studies that found MDMA altered speech when participants were instructed to describe a loved one (Baggott et al., 2015; Bedi et al., 2014; Wardle and de Wit, 2014). However, only one previous study examined if MDMA altered experience of specific autobiographical memories. Carhart-Harris et al. (2014) found that participants cued to remember positive memories after MDMA rated them as significantly more positive, vivid, and emotionally intense, while worst memories were rated as less negative, compared to after placebo. We hypothesized that autobiographical descriptions of positively and negatively valenced memories might reveal MDMA-induced changes in processing of autobiographical memories including participants feeling increased comfort and insight when recounting these events.

## Methods

### General Study Design

We used a double-blind, placebo-controlled, within-subject, gender-balanced design. In two experimental sessions that were separated by at least one week, twelve volunteers (6 male, 6 female) experienced placebo and 1.5 mg/kg oral MDMA after an overnight hospital stay. Participants were discharged 6 h after MDMA or placebo or after drug effects resolved, whichever was later, and they returned at 24 hours for a brief visit. We selected 1.5 mg/kg MDMA, measured as the hydrochloride salt, as an active dose to would produce typical drug effects based on past clinical studies (e.g., (Lester et al., 2000; Tancer and Johanson, 2001; Cami et al., 2000; Vollenweider et al., 1998; Harris et al., 2002)). To ensure adequate blinding, participants consented to take 1 or 2 active doses of MDMA, even though all participants received 1 active dose and 1 placebo. This study also included other measures, including physiological measures of effects of MDMA on hydration status, which are being prepared for separate publication and are not described here.

### Participants

We recruited healthy, MDMA-experienced individuals between the ages of 18 and 50, through newspaper and on-line advertisements and word-of-mouth. A licensed physician determined participants to be healthy based on medical questionnaires, laboratory screenings, and a physical exam. Exclusion criteria included DSM-IV dependence on MDMA or any other psychoactive drug (except nicotine or caffeine); desire to quit or decrease MDMA use; history of adverse reaction to study drugs; current enrollment in a drug treatment program; current supervision by the legal system; any current physical or psychiatric illness that might be complicated by the study drugs or impair ability to complete the study; body mass index (weight/height^2^) greater than 30 or less than 18 kg/m^2^; and current or recent use of any medication that might pose risk of drug-drug interaction.

## Experimental measures

### Autobiographical memory task

We developed a novel task to measure MDMA effects on autobiographical memory recall. During drug administration sessions, beginning 1.5 - 2 hours after MDMA/placebo, participants were given 5 minutes per memory to describe autobiographical memories from each of four emotional categories and a “typical day” (i.e., participants spoke for 25 min total). Emotional categories were: fear (defined as “afraid, terrified, or extremely anxious”); safe (“safe, comfortable, secure, or protected”); sad (“sad, at a loss, mournful, or depressed”); and joy (“feel joyful, happy, ecstatic, or in love with life”). We selected these to include both high and low arousal events with positive and negative valences. After recounting each event, participants rated their mood and experience of describing the memory. The order for these autobiographical memories was randomized with the exception that Joy was always described last, in order to minimize any residual discomfort. A researcher was present and listened but was largely silent, except for answering direct questions.

### Selecting comparable autobiographical events

To ensure autobiographical events were well balanced between conditions, we collected lists of candidate experiences in a separate screening session. Participants were asked to remember several of the most powerful non-drug-related experiences they could for each category. Participants rated candidate episodes using the intrusion scale of the Impact of Life Events Scale (Horowitz et al., 1979), the Positive and Negative Affect Schedule (PANAS) (Watson et al., 1988), and a series of questions on event recency, vividness of sensory/perceptual detail, level of personal involvement (passive/bystander vs. active), level of consequence to self, confidence in accuracy (See **Table 1** for details). We then selected two comparable episodes for each emotional category and randomly assigned them to the two sessions.

**Table 1:**
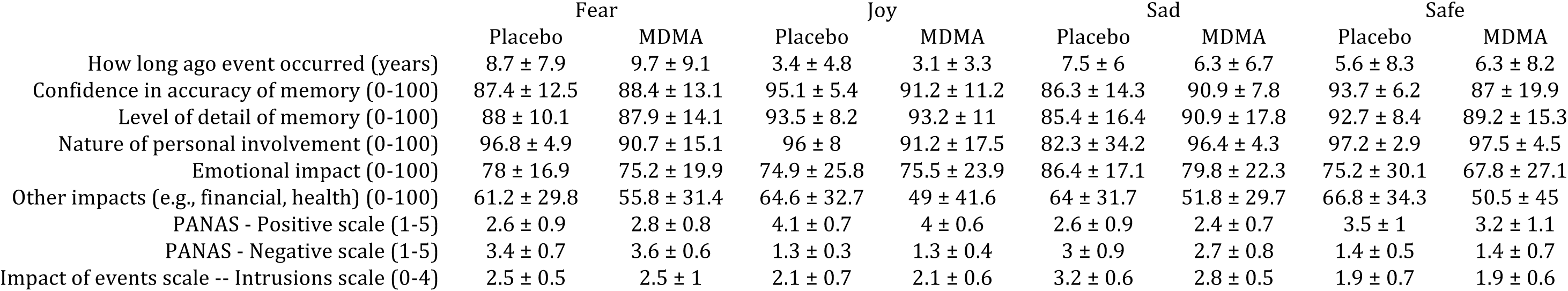
Autobiographical memories used in placebo and MDMA conditions were comparable in multiple dimensions. Values are given as mean ± SEM.

### Experience of remembering measures

After describing each episode, participants made the following visual analog ratings: How upsetting was it for you to talk about this experience (hereafter, upsetting talking); How comfortable was it for you to talk about this experience (comfortable talking); How much did you re-live the emotions you felt when you had this experience (relive emotions); Rate your ability to remember the details of the experience (ability to remember details); Rate your ability to remember the emotions you felt during this experience (ability to remember emotions); Rate your ability to describe the emotions you felt during this experience (ability to describe emotions); and Rate your ability to understand the emotions you felt during this experience (ability to understand emotions). Questions beginning with “how” used the anchors “not at all” and “completely”; those beginning with “rate” used the anchors “much worse than usual” and “much better than usual.” These ratings were analyzed using mixed-effects models with participant as a random effect and fixed effects for emotion category, drug condition, and the interaction of the two.

### Analysis of narratives

We transcribed autobiographical memories and analyzed them using Pennebaker’s Linguistic Inquiry and Word Count 2007 (LIWC, version 1.11), which has been used extensively to analyze speech and text samples in past studies (reviewed in Tausczik and Pennebaker 2010). LIWC is a word count program that matches text against an extensive dictionary, and provides the percent of words in a large set of well-validated categories. We examined the same 43 categories we had used in Baggott et al. (2015). Reliability and validity information has been reported by Pennebaker and King (1999).

## Self-report measures

### Visual analog items

We measured the time course of self-report drug effects using visual analog items intended to tap general drug effects (Any Drug Effect, Drug Liking, Good Drug Effect, High), stimulant/sedative effects (Anxious, Clear headed, Confused, Drunken, Relaxed, Stimulated), and other emotional effects (Adventurous, Amused, Closeness to others, Contented, Enthusiastic, Insightful, Kind, Loving, Passionate, Proud, Trusting). These were collected before and 1, 2, 3, 4, 5, and 6 hours after drug administration. Participants clicked a location on a digital line to indicate how intensely they were experiencing each of these items in the last few minutes. Responses were scored on a scale of 0 to 100. Peak baseline-subtracted responses were used in statistical models.

### Social anxiety

We measured social anxiety using the Brief Fear of Negative Evaluation – revised (BFNE) (Carleton et al., 2006; Leary, 1983). This 12-item Likert scale questionnaire measures apprehension and distress due to concerns about being judged disparagingly or with hostility by others. It is believed that this is a fundamental fear distinct from concern about illness or injury (Taylor, 1993). Pilot testing indicated that our healthy MDMA-experienced participants tended to give very low ratings on this measure, limiting sensitivity to potential decreases. Therefore, we modified the instrument to use a 5-point Likert scale with the lowest, middle, and highest values labeled with “Much less than normal”, “normal”, and “Much more than normal”. The BFNE was administered before and 1.5, 2, 2.5, and 24 h after drug administration, shortly after the autobiographical memory task was completed. Participants were instructed to answer for how they had been feeling for the past hour. Baseline-subtracted responses were used in statistical models.

### Authenticity

We measured MDMA effects using the 45-item Authenticity Index (Kernis and Goldman, 2006), which seeks to measure feelings of ‘‘unimpeded operation of one’s true- or core-self.” Because the Authenticity Index was designed to measure authenticity as a trait, we slightly modified the instructions and some items to measure current feelings. The typical change was that items describing usual behaviors were modified to refer to the present or to hypotheticals situations (original and modified items are included as supplemental material. We used the same Likert scale and anchors as with the BFNE. The Authenticity Index was originally reported to have a total score made up of four subscales. However, White (2011) was unable to replicate the factor structure of the instrument. Therefore, we are only reporting total scores. The authenticity index was given at 2.5 h after drug administration, shortly after the autobiographical memory task was completed. Participants were instructed to answer for how they had been feeling for the past hour.

### Interpersonal functioning

We measured interpersonal functioning using the Interpersonal Adjectives Scale-Revised (IASR) (Trapnell and Wiggins, 1990; Wiggins et al., 1988). The IASR is a widely used self-report measure of interpersonal functioning in which 8 subscales or octants are evenly distributed as vectors originating at the origin of a two dimensional space that can be labeled as Dominance or Agency (concern for mastery and power that enhance and protect the individual) on the vertical axis and Nurturance or Communion (a concern for intimacy and solidarity with others) on the horizontal axis (Kiesler, 1991; Wiggins and Broughton, 1991). The two dimensions of Dominance and Nurturance can be considered as a 45-degree rotation of the Big Five personality dimensions Extraversion and Agreeableness (McCrae and Costa, 1989). To reduce the duration of the instrument, we used the 32 highest loading items from the original 64-item instrument, as in Knutson (1996). The IASR was given at 2.5 hours after drug administration, shortly after the autobiographical memory task was completed. Participants were instructed to answer for how they had been feeling for the past hour.

## Results

### Participant characteristics

Twelve participants (6 male, 6 female), ages 29 ± 2 years (mean ± SEM, range: 21-40) with 24 ± 7 (range 5-75) previous MDMA experiences, completed the study.

### Autobiographical memory task

The autobiographical memories used in the study did not significantly differ between the two conditions based on measures of their recency and impact, as summarized in **Table 1**. Events generally involved personal relationships, achievements/failures, family events, travel, accidents/illnesses, sexual violence, and property damage.

#### Experience of remembering

Participants reported feeling more comfortable talking about emotional memories while on MDMA, shown in **Figure 1**. For ratings of how comfortable participants felt talking, a mixed effects model with a random effect of participant revealed significant fixed effects of condition (F_1,80_ = 5.30, p = 0.024) and emotion (F_3,80_ = 3.77, p = 0.014). Participants reported feeling 10.4 ± 4.5 visual analog units more comfortable on MDMA compared to placebo. In addition, memories of Joy were significantly easier to recount than other emotions (p = 0.006, 0.005, and 0.015 comparing Joy to Sad, Fear, and Safe, respectively).

**Figure 1:**
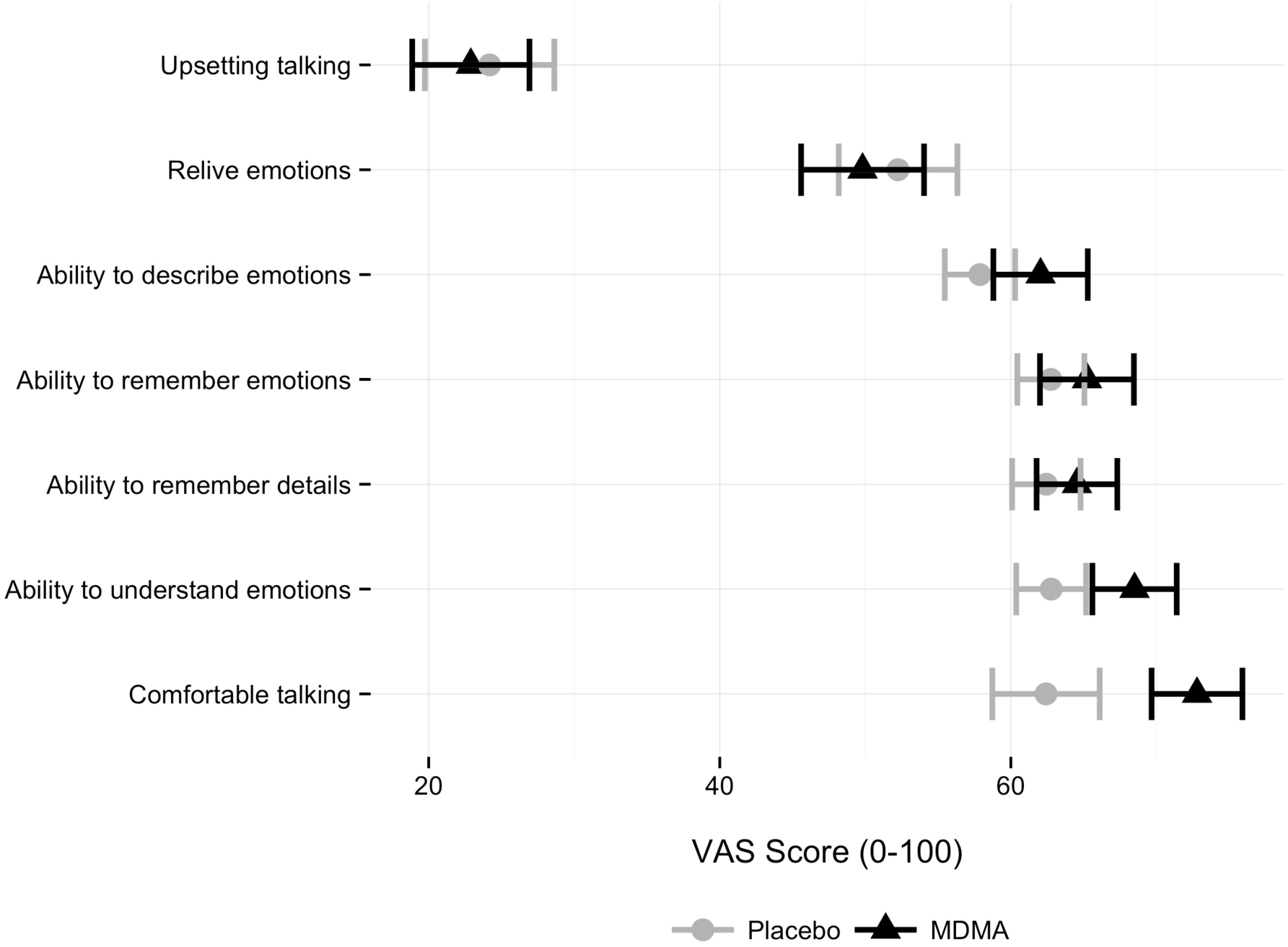
MDMA increased participants’ comfort describing emotional memories. Drug effects on participant ratings of the experience describing autobiographical memories. Placebo is gray circles, MDMA is black triangles.

MDMA did not otherwise appear to alter significantly participants’ reports of their abilities to remember, understand, or experience their emotional memories. However, there was a trend for participants to report greater understanding of their memories on MDMA compared to placebo (p = 0.058). The emotional category of the memory was a significant predictor of several aspects of remembering. Specifically, there were main effects of emotion for reliving the experience (F_3,77_ = 3.47, p = 0.020) and feeling upset talking about the experience (F_3,77_ = 11.8, p < 0.001) and there were trends for an effect of emotion on ability to remember details (p = 0.070), remember emotions (p = 0.088), and describe emotions (p = 0.075).

### Analysis of narratives

MDMA decreased the number of spoken words and altered word choice in several categories. MDMA significantly decreased the number of words participants spoke (F_1,107_ = 9.46, p = 0.003) from 651.8 ± 42.33 words after placebo to 592.3 ± 45.98 words after MDMA. For each LIWC category, we fit a mixed effects model predicting LIWC scores using participant as a random effect and including fixed effects for condition and emotional memory category and their interaction. MDMA increased use of *present tense* (F_1,77_ = 8.22, p = 0.005), words showing *assent* (e.g., agree, okay, yes; F_1,77_ = 5.07, p = 0.027), and words relating to *family* (e.g., daughter, husband, aunt; F_1,77_ = 4.51, p = 0.037). We detected a non-significant trend for MDMA to increase use of words in the larger *social* category (e.g., talk, they, child; p = 0.099) of which *family* is a subcategory. There were also non-significant trends for MDMA to increase words relating to *finance* (e.g., audit, cash, owe; p = 0.059) and to decrease those relating to *anger* (e.g., hate, kill, annoyed; p = 0.095).

Fourteen of the 43 LIWC categories showed significant effects of category of emotional memory being recalled. LIWC categories that varied based on emotional condition largely related to emotional and social language (LIWC categories: *affective processes*, *positive emotion*, *negative emotion*, *anger*, *anxiety*, *sad*, *feel*, *social processes*), with five more general categories also showing effects (*leisure, death, religion, space, home*). F-values for the main effect of emotional memory category in these models ranged from 2.86 to 50.1 (with 1 and 77 degrees of freedom), while p-values were from 0.042 to less than 0.001.

### Self-report measures

Self-report data were missing eight VAS items for one participant at one placebo time point due to a computer failure. The VAS item Self-conscious was missing at all times for two participants after placebo and one participant after MDMA due to a version control error.

#### Visual analog measures

When visual analog measures were examined, we found MDMA produced robust increases in measures of general drug effects, stimulant- and sedative-like effects, and changes in emotional measures of love and kindness, as shown in **Figure 2** and **Table 2.** In linear mixed effects models with drug condition as a fixed effect and participant as a random effect, MDMA condition predicted peak increases in Any Drug Effect (F_1,11_ = 82.35, p < 0.001), Good Drug Effect (F_1,11_ = 103.18, p < 0.001), Drug Liking (F_1,11_ = 95.12, p < 0.001), and High (F_1,11_ = 82.08, p < 0.001).

**Table 2:** MDMA had robust effects on peak visual analog measures of general drug effects and stimulant/sedative effects, while having less consistent effects on emotional items. Placebo is gray circles, MDMA is black triangles. Values are given as mean ± SEM. The symbols >, <, and indicate greater than, less than, and not significantly different.

|  |  | Placebo |  | MDMA | Effect of Condition |
| --- | --- | --- | --- | --- | --- |
| General Effects | Any Drug Effect | 19.6 ± 8.8 | < | 88.5 ± 3.0 | F1,11 = 82.35, p < 0.001 |
|  | Good Drug Effect | 18.3 ± 8.4 | < | 86.2 ± 2.9 | F1,11 = 103.18, p < 0.001 |
|  | High | 16.4 ± 8.5 | < | 85.3 ± 2.5 | F1,11 = 82.08, p < 0.001 |
|  | Drug Liking | 18.0 ± 9.0 | < | 85.1 ± 3.5 | F1,11 = 95.12, p < 0.001 |
| Emotional Effects | Adventurous | 11.8 ± 3.4 | ≈ | 16.9 ± 5.4 | F1,10 = 1.91, p = 0.197, ns |
|  | Amused | 25.9 ± 6.9 | ≈ | 26.9 ± 4.7 | F1,10 = 0.01, p = 0.910, ns |
|  | Closeness to others | 24.1 ± 4.9 | ≈ | 35.5 ± 6.4 | F1,11 = 2.05, p = 0.180, ns |
|  | Content | 11.3 ± 3.1 | ≈ | 17.8 ± 3.3 | F1,10 = 3.21, p = 0.103, ns |
|  | Enthusiastic | 16.7 ± 5.2 | < | 34.4 ± 5.6 | F1,10 = 5.38, p = 0.043 |
|  | Insightful | 21.4 ± 6.5 | ≈ | 28.7 ± 8.5 | F1,11 = 0.55, p = 0.475, ns |
|  | Kind | 11.7 ± 3.3 | < | 25.7 ± 4.1 | F1,10 = 7.24, p = 0.023 |
|  | Loving | 12.3 ± 3.6 | < | 26.3 ± 5.4 | F1,10 = 6.53, p = 0.029 |
|  | Passionate | 16.7 ± 4.2 | ≈ | 26.6 ± 6.1 | F1,10 = 2.35, p = 0.156, ns |
|  | Proud | 12.9 ± 2.8 | ≈ | 21.4 ± 5.6 | F1,10 = 1.95, p = 0.193, ns |
|  | Self-conscious | 21.7 ± 6.8 | ≈ | 24.4 ± 6.4 | F1,7 = 0.08, p = 0.782, ns |
|  | Trusting | 7.8 ± 2.0 | ≈ | 12.4 ± 4.5 | F1,11 = 1.23, p = 0.291, ns |
| Stimulant/Sedative | Anxious | 16.1 ± 4.6 | < | 41.6 ± 7.1 | F1,11 = 9.49, p = 0.010 |
|  | Clear headed | 4.1 ± 1.9 | ≈ | 14.3 ± 5.6 | F1,11 = 3.43, p = 0.091 |
|  | Confused | 15.0 ± 5.6 | ≈ | 22.9 ± 6.2 | F1,11 = 0.95, p = 0.350, ns |
|  | Drunken | 3.3 ± 2.5 | < | 41.8 ± 10.07 | F1,11 = 18.26, p = 0.001 |
|  | Relaxed | -11.1 ± 3.8 | > | -33.3 ± 5.5 | F1,11 = 11.05, p = 0.007 |
|  | Stimulated | 24.6 ± 7.5 | < | 61.2 ± 7.4 | F1,11 = 21.98, p < 0.001 |

**Figure 2:**
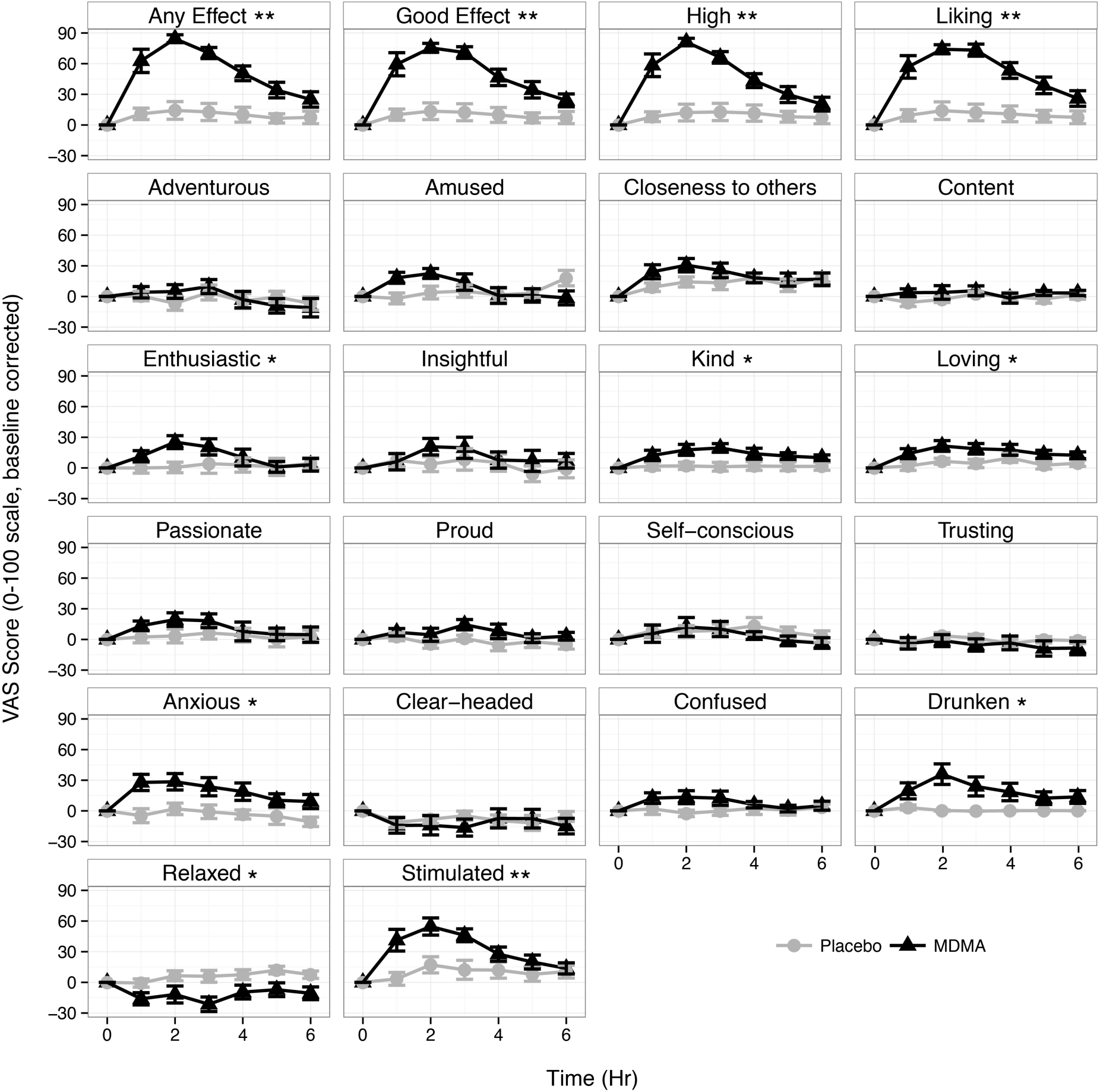
Visual analog measures of the time course of drug effects. Measures are sorted based on whether they are general drug effects (top row), putative emotional effects (middle three rows), or stimulant/sedative effects (bottom two rows). Significant drug effects on maximum absolute change from baseline are indicated with * for p < 0.05 and ** for p < 0.001.

MDMA increased ratings of both stimulant- and sedative-like feelings, including peak increases in Anxious (F_1,11_ = 9.49, p = 0.010), Drunken (F_1,11_ = 18.26, p = 0.001), Enthusiastic (F_1,10_ = 5.38, p = 0.043), and Stimulated (F1,11 = 21.98, p < 0.001) ratings along with decreases in ratings of Relaxed (F_1,11_ = 9.46, p = 0.011). There was also a trend for participants to feel less Clear-headed on MDMA (p = 0.091).

MDMA increased feelings of love and kindness. In individual mixed effects models with drug condition as a fixed effect and participant as a random effect, there were main effects of condition on peak Loving (F_1,10_ = 6.53, p = 0.029) and Kind (F_1,10_ = 7.24, p = 0.023) ratings. Scores for Closeness To Others in both conditions appeared to show an elevation at 6 hours associated with discharge from the hospital. When that time-point was excluded, there was a trend for an effect of condition (p = 0.09). MDMA did not significantly affect peak ratings of Adventurous, Amused, Contented, Insightful, Proud, Passionate, Self-conscious, or Trusting.

#### Social anxiety

MDMA decreased social anxiety, as shown in **Figure 3A**. In analogous mixed effects models to those previous used, MDMA decreased maximum magnitude change from baseline BFNE scores (F_1,10_ = 7.7, p = 0.019). An (underpowered) exploratory analysis that included gender and a condition by gender interaction term did not reveal a significant effect of gender.

**Figure 3:**
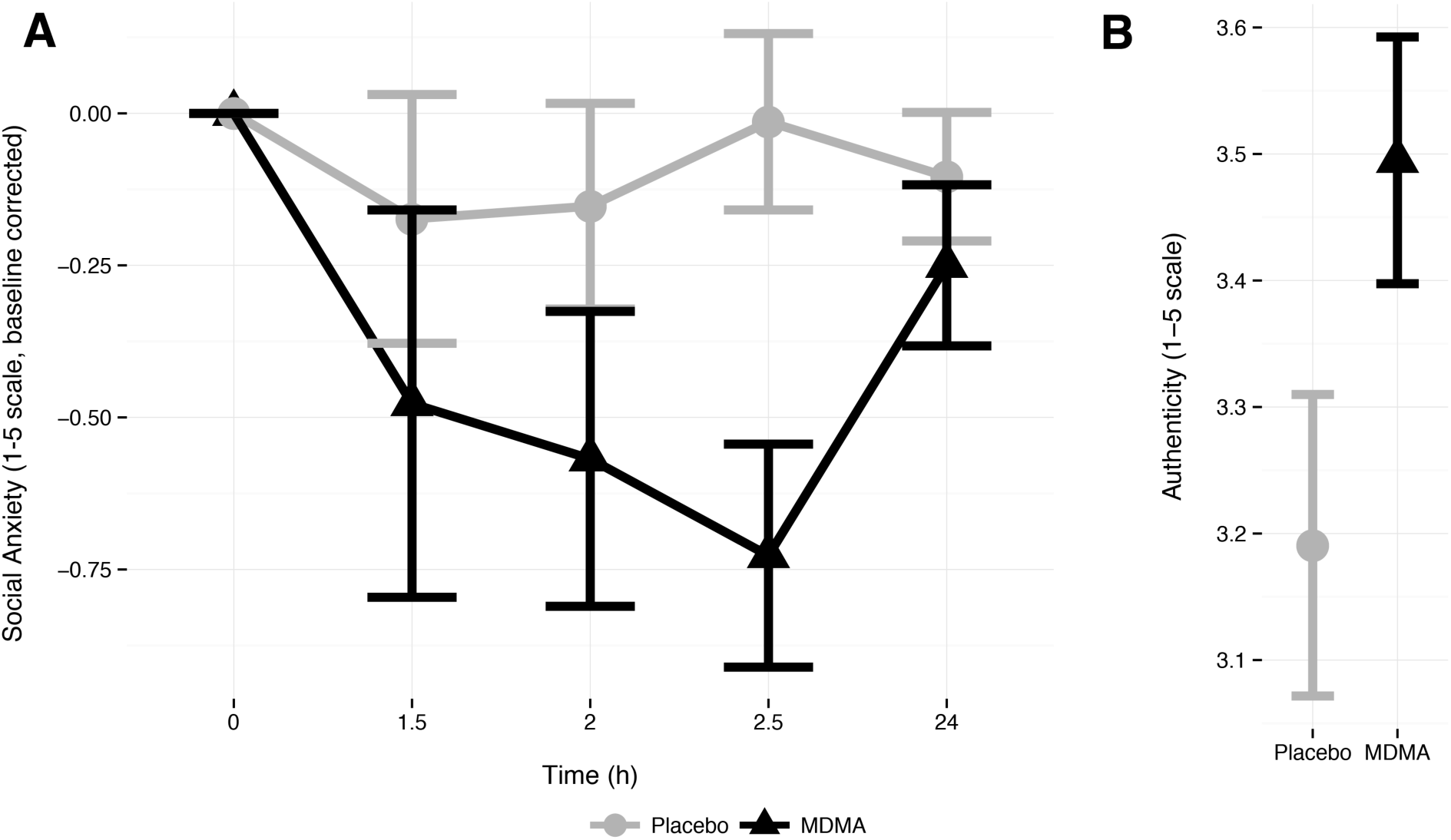
MDMA decreased self-report social anxiety (A) and increased feelings of authenticity (B). Social anxiety was measured with the BFNE, while authenticity was measured with the Authenticity Index. Placebo is gray circles, MDMA is black triangles.

#### Authenticity

MDMA increased feelings of authenticity, as shown in **Figure 3B**. There was a main effect of condition F_1,10_ = 12.07, p= 0.006) on total authenticity score. Exploratory addition of gender and gender by condition terms did not reveal significant effects.

#### Interpersonal functioning

As shown in **Figure 4**, MDMA increased affiliative feelings, measured as an increase (right shift) in the Nurturance/Communion dimension. This shift appeared to be mainly caused by significant increases in the Gregarious subscale. In a mixed effects model predicting Nurturance with participant as a random effect and condition as a fixed effect, there was a significant effect of condition (F_1,11_ = 5.52, p = 0.039). In analogous models of the individual subscales, condition predicted an increase in the Gregarious (F_1,11_ = 8.49, p = 0.014) subscale and there were statistical trends for an increase in the Assured subscale (p = 0.077) and a decrease in the Aloof (p = 0.060) subscales. No significant changes were detected in the Dominance dimension.

**Figure 4:**
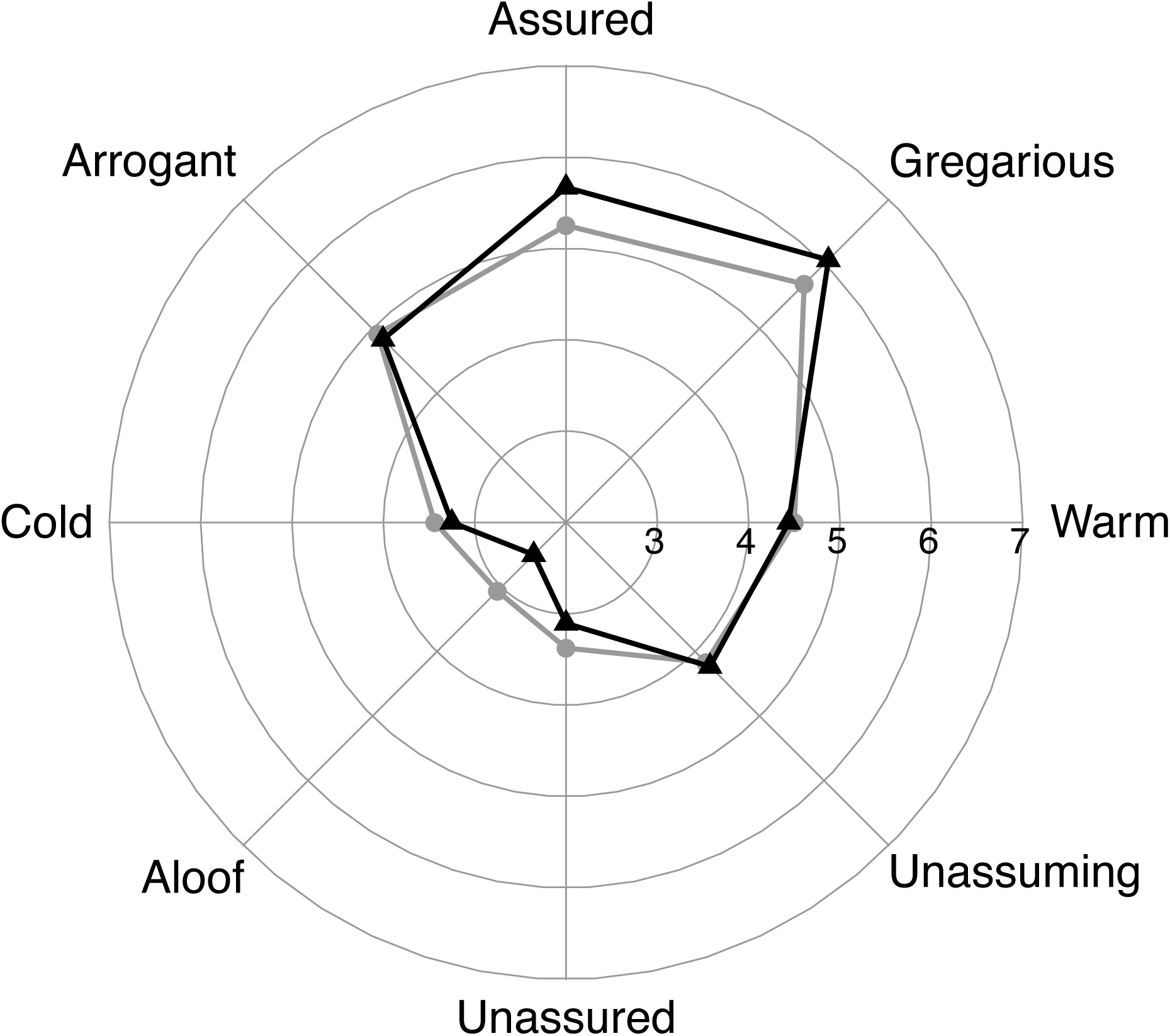
MDMA made participants feel more Gregarious. The IASR measures self-report interpersonal functioning questionnaire using a circumplex model in which eight evenly spaced scales sample aspects of interpersonal functioning within a two dimensional space where the vertical dimension indicates Dominance and the horizontal indicates Nurturance/Communion. MDMA increased the Gregarious scale and shifted the overall “center of mass” to the right, an increase in the Nurturance/Communion dimension.

### Discussion

We conducted a double-blind placebo controlled study of the effects of MDMA on social emotions and autobiographical disclosure in a controlled setting. We found that MDMA simultaneously positively altered evaluation of the self (i.e., increasing feelings of authenticity) while decreasing concerns about negative evaluation by others (i.e., decreasing social anxiety). Consistent with these feelings, MDMA increased how comfortable participants felt describing emotional memories. Overall, MDMA produced a prosocial syndrome that seemed to facilitate emotional disclosure.

MDMA is sometimes described as decreasing fear (e.g., Greer and Tolbert, 1990). However, our study instead suggests a more focused effect on social anxiety rather than a general anxiolytic effect. We found that MDMA increased self-report anxiety, as had been previously seen in some studies, but also decreased social anxiety, which is a novel finding. This pattern is inconsistent with the idea that MDMA is a general anxiolytic. However, it is consistent with the possibility that MDMA may facilitate an affiliative “tend-and-befriend” style of response to stressors. This alternative to the prototypical male “fight or flight” response involves stress-induced caregiving and prosocial behavior (Taylor, 2006). The response to MDMA in our study — participants reported feeling more anxious yet also more affiliative, kind, and loving — appears reminiscent of a tend-and-befriend response. Moreover, both the tend-and-befriend response and MDMA have been proposed to share a common neural mechanism of oxytocin release (Taylor, 2006; Thompson et al., 2007). Whether or not this hypothesis proves accurate, our findings are consistent with the theory that MDMA could aid psychotherapy by improving the therapeutic alliance, particularly when dealing with stressful autobiographical material.

Participants reported feeling greater comfort disclosing emotional autobiographical episodes after MDMA compared to after placebo. We did not detect other effects of MDMA on remembering, describing, or understanding emotional memories. We had hypothesized that participants would feel more insightful and report greater understanding of their memories. However, we could not confirm this. This may be because these effects were absent, the memories had been recently recalled in a screening session and were already well understood, or because we were underpowered to detect them given our modest sample size.

Similarly, we detected changes in speech that only partly overlapped with those seen in past studies. Wardle and de Wit (2014) found MDMA increased positive word use in a speech task in which participants described a loved one. Baggott et al. (2015), using the same task, found that MDMA caused participants to use more words in categories relating to social processes, sexuality, and death. We did see changes in a social subcategory relating to family and saw a trend for changes in the main social category. However, we did not replicate the specific findings relating to positive emotion, sexuality, or death. These differences may be the result of task differences (describing a loved one vs. describing emotional memories), relative lack of power in the current study, or Type II errors. More generally, these results and those using less comparable machine learning analyses (Bedi et al., 2014; Baggott et al., 2015) seem to point to greater emphasis on social topics and an increased willingness to disclose after MDMA.

Participants reported feeling greater authenticity after MDMA. Authenticity is an idea that has its roots in humanistic psychology and refers to the unimpeded operation of one’s true self (Maslow, 1968; Rogers, 1961). This feeling that one is able to be oneself and can reduce self-censorship is associated with greater well-being, more honesty, and lessened defensiveness (Maltby et al., 2012; Kernis and Goldman, 2006; Lakey et al., 2008; Wood et al., 2008). Consistent feelings of authenticity seem likely to be an effect of MDMA that distinguishes it from classical psychedelics such as LSD and psilocybin. Psychedelics may produce feelings of insight into one’s true self yet they also often produce depersonalization, the feeling that one is not oneself (Studerus et al., 2011; Hollister, 1968). It remains to be seen to what extent authenticity distinguishes MDMA from stimulants such as amphetamine, especially since state authenticity can be increased by positive mood (Lenton et al., 2013).

This study had several limitations. The sample size was modest for studies of psychological drug effects and we were likely underpowered to detect less-than-robust effects. Autobiographical memory narratives were truncated when they extended beyond 5 minutes, which may have hampered ability to detect drug effects if salient features were not evenly distributed throughout narratives. We attempted to match autobiographical memories, but in retrospect failed to control for temporal duration (and resulting narrative complexity) of memories. In addition, memories of feeling “safe,” which we hoped would reflect a positive low arousal state, often included descriptions of initial fear and danger. We modified the anchors of some questionnaires (BFNE, Authenticity Index) and reworded trait items to reflect state in the Authenticity Index. Thus, there would be value in measuring MDMA effects with other measures of social anxiety and authenticity. Finally, the current study did not assess whether any of the measured effects of MDMA were unusual to that drug as could be done by comparing it to a stimulant like methamphetamine, which was once suggested to aid psychotherapy with reasoning reminiscent to that used for MDMA (e.g., Levine et al., 1948; Ling and Davies, 1952).

In our view, research on MDMA continues to be challenged by the difficulties of reliably measuring the unusual effects of MDMA. General measures of drug effects, such as the VAS item Good Drug Effect, are exquisitely sensitive to MDMA but tell us little about the socioemotional specifics. Many socioemotional measures, such as categorizing or rating emotional stimuli, yield subtle effects and thus appear to be relatively insensitive to the robust MDMA syndrome. Other measures that are more specific, such as the VAS item “Love”, have high variance and ceiling effects and are sensitive to interpersonal context, which is often impoverished in a psychopharmacological setting. In this study, we attempted to address these issues both by creating a controlled setting reminiscent of psychotherapy and by adapting several socioemotional outcome measures. We found that MDMA decreased social anxiety, increased sociability and feelings of authenticity, and enhanced comfort disclosing autobiographical material. These effects occurred against a background in which MDMA had both stimulant-like (e.g., stimulation and anxiety) and sedative-like (e.g., VAS item indicating feeling drunk) self-report effects. Although conclusive studies are lacking, MDMA appears to have unusual socioemotional effects, consistent with the proposal that it represents a new class of psychoactive with psychotherapeutic potential (Nichols, 1986).

## Acknowledgements

This research was supported by National Institutes of Health DA017716 and DA016776 and the NIH/National Center for Research Resources UCSF-CTSI UL1 RR024131. The authors thank Ryne Didier, Margie Jang, and Juan Carlos Lopez for assistance in the study and Josh Tetrick for his support.

